# ARHGEF18/p114RhoGEF coordinates PKA/CREB signaling and actomyosin remodeling to drive trophoblast cell-cell fusion during placenta morphogenesis

**DOI:** 10.1101/2020.07.12.199141

**Authors:** Robert Beal, Ana Alonso-Carriazo Fernandez, Dimitris K. Grammatopoulos, Karl Matter, Maria S. Balda

**Author notes:** Address for correspondence UCL Institute of Ophthalmology, University College London, Bath Street, London EC1V 9EL, UK, /6861, /. lead contact Maria S. Balda.

## Abstract

Coordination of cell-cell adhesion, actomyosin dynamics and gene expression is crucial for morphogenetic processes underlying tissue and organ development. Rho GTPases are main regulators of the cytoskeleton and adhesion. They are activated by guanine nucleotide exchange factors in a spatially and temporally controlled manner. However, the roles of these Rho GTPase activators during complex developmental processes are still poorly understood. ARHGEF18/p114RhoGEF is a tight junction-associated RhoA activator that forms complexes with myosin II, and regulates actomyosin contractility. Here we show that p114RhoGEF/ ARHGEF18 is required for mouse syncytiotrophoblast differentiation and placenta development. *In vitro* and *in vivo* experiments identify that p114RhoGEF controls expression of AKAP12, a protein regulating PKA signalling, and is required for PKA-induced actomyosin remodelling, CREB-driven gene expression of proteins required for trophoblast differentiation, and, hence, trophoblast cell-cell fusion. Our data thus indicate that p114RhoGEF links actomyosin dynamics and cell-cell junctions to PKA/CREB signalling, gene expression and cell-cell fusion.

## INTRODUCTION

Development of tissues and organs is mediated by complex morphogenetic processes that require remodeling of cell-substrate and cell-cell adhesion, gene expression, as well as dynamic cellular processes driven by the actomyosin cytoskeleton. Rho GTPases are major regulators of these processes. They regulate a wide variety of different cellular mechanisms but signal via a relatively small set of effector molecules; hence, it is thought that the molecular mechanisms that control their activation and inactivation in space and time determine process-specificity. Activation is mediated by guanine nucleotide exchange factors (GEFs) that catalyze the exchange of GDP by GTP. A large number of GEFs forming two protein families, Dbl and DOCK GEFs, have been identified and characterized in various *in vitro* systems; however, their functions in tissue and organ morphogenesis, and their interactions with major signalling pathways that drive gene expression and cell differentiation are not well understood.

Cell-cell adhesion complexes such as tight and adherens junctions interact with the cytoskeleton and harbor regulatory proteins that control cytoskeletal dynamics and, thereby, cell adhesion and behavior. A key group of such signaling proteins recruited to tight junctions are GEFs for RhoA, which includes p114RhoGEF/ARHGEF18, GEF-H1/ARHGEF2 and ARHGEF11 (Benais-Pont et al., 2003; Itoh et al., 2012; Kim et al., 2015; Terry et al., 2011; Xu et al., 2013). However, their roles in developmental morphogenetic processes are still poorly understood. Mutations in fish indicate that p114RhoGEF may function in the maintenance, rather than development, of apicobasal polarity in neuroepithelia (Herder et al., 2013). In humans, partially inactivating p114RhoGEF mutations lead to retinitis pigmentosa after apparently normal retinal development (Arno et al., 2017). A genome-wide SNP analysis also linked p114RhoGEF to capillary leak syndrome (Clarkson disease); however, the effects of the SNPs on p114RhoGEF activity, and the underlying molecular and cellular processes linking p114RhoGEF to vascular leakage are not known (Xie et al., 2013). Hence, the roles of p114RhoGEF in cell adhesion dynamics in vivo and tissue morphogenesis are not known.

Given the role of p114RhoGEF in the regulation of dynamic cellular processes and the coordination of actomyosin activation in response to changes in cell adhesion *in vitro* (Acharya et al., 2018; Haas et al., 2020; Nakajima and Tanoue, 2011; Terry et al., 2012; Terry et al., 2011; Zihni et al., 2017), we asked whether such functions are important for organ morphogenesis. Our data show that p114RhoGEF is indeed essential for mouse development with embryos displaying a number of different phenotypes. A main phenotype observed in p114RhoGEF-deficient mice is the failure of normal placenta development, a PKA-driven process that involves cell-cell fusion during syncytiotrophoblast formation, which requires activation of PKA/CREB-induced expression of the transcription factor Gcm1 and proteins that act as fusogens, syncytins, as well as remodeling of the actin cytoskeleton. Results from knockout mice and cultured trophoblast models indeed indicate that p114RhoGEF is drives for cell-cell fusion by coordinating actomyosin remodeling and PKA/CREB-regulated expression of proteins required for cell-cell fusion.

## RESULTS

### p114RhoGEF is required for mouse and placenta development

We first tested whether p114RhoGEF is important for mammalian embryonic development in a global p114RhoGEF knockout mouse strain (p114^−/−^) from EMMA (ID EM:02310) that carries a retroviral gene trap insertion in the intron before the first coding exon (Fig. 1A). Immunoblotting of embryos at E12.5 revealed that the insertion resulted in efficient knockout of p114RhoGEF protein expression (Fig. 1B). Analysis of born litters from mating heterozygous mice revealed that no p114^−/−^ mice were born (Supplementary Fig. 1A), indicating embryonic lethality. During embryonic development, the expected numbers of p114^−/−^ embryos were present at E11.5 and E12.5 (Fig. 1C), but by E15.5 the number of p114^−/−^ embryos was strongly reduced. More detailed analysis of the phenotypes of p114^−/−^ embryos at E10.5/11.5 and E12.5/13.5 revealed various phenotypes including anemia, abdominal hemorrhages and effusions as well as yolk sac edema and vascular defects (Fig. 1D,E; Supplementary Fig. 1B,C1). At E12.5, a large number of embryos had placenta defects and/or exhibited growth arrest or resorption (Fig. 1D,E). Placenta defects are a common cause of embryonic lethality in mouse mutants because of its importance in maternal-fetal nutrient exchange (Perez-Garcia et al., 2018). Hence, defects in placenta development - although they may not be the only reason for embryonic death - are likely to make an important contribution to the lethality of p114RhoGEF-deficient embryos.

**Figure 1:**
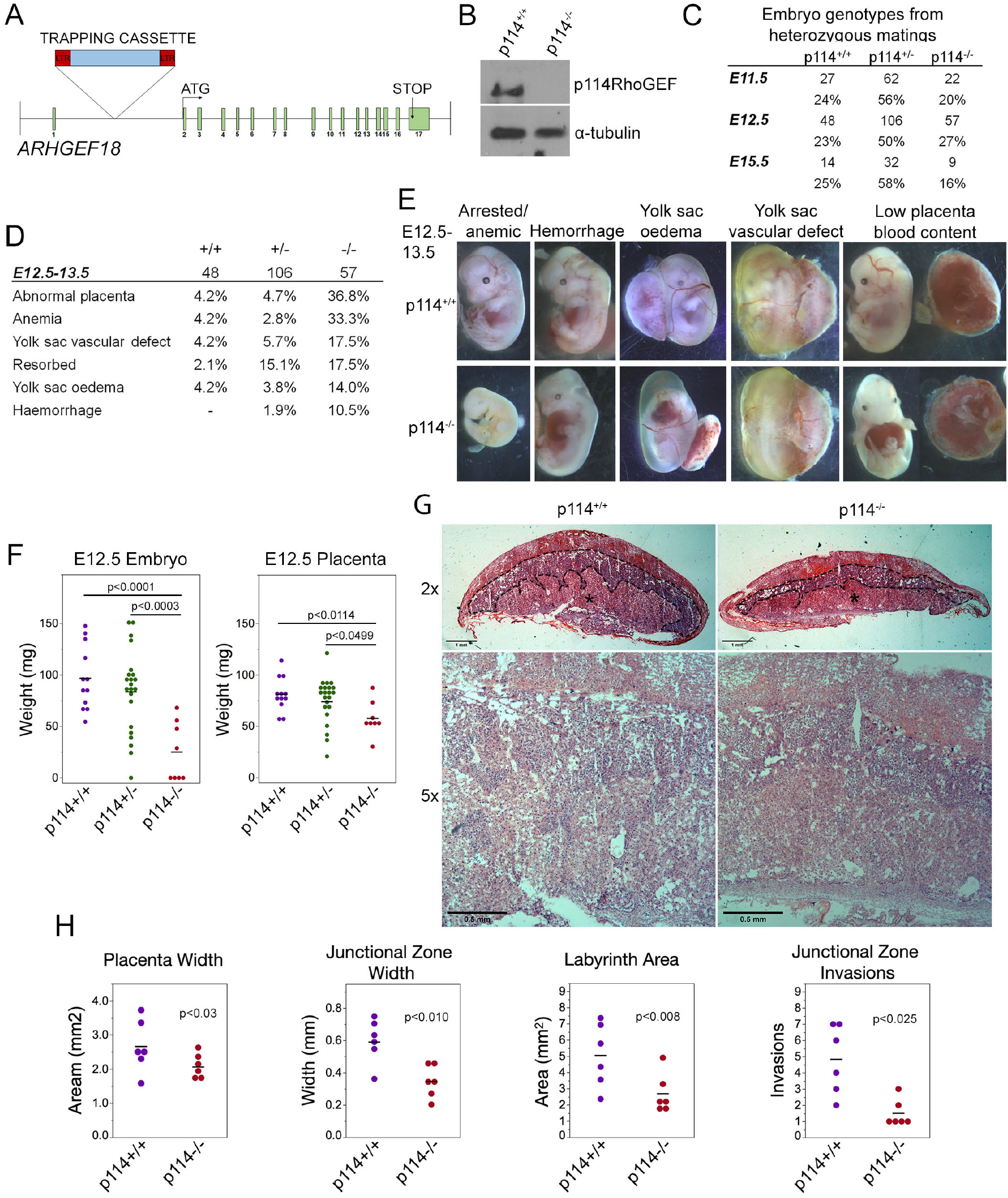
Knockout of p114RhoGEF in mice is embryonic lethal. (**A**) Schematic representation of mutation strategy for generating the ARGEF18/p114 KO mouse (adapted from https://www.taconic.com/knockout-mouse/arhgef18-trapped). Indicated is the insertion site of the trapping cassette prior to the start initiation codon. (**B**) Immunoblot of samples generated from E12.5 embryos, indicating complete ablation of p114RhoGEF expression in p114^−/−^ embryos. (**C**) Analysis of embryos at E11.5, E12.5 and E15.5. Genotypes at E12.5 were at expected mendelian ratios but by E15.5, the percentage of p114^−/−^ embryos started to decrease due to embryonic arrest. (**D,E**) Analysis of p114^−/−^ phenotypes at E11.5 and representative images of phenotypes observed. (**F**) Analysis of embryo and placenta weights at E12.5. Shown are data points and means. Significance was tested with t-tests. **(G)** Hematoxylin and eosin (H&E) staining of 12μm placenta cryosections at E12.5. Note the decrease in thickness of the labyrinth layer in p114RhoGEF-deficient animals (asterisks). Junctional zones are indicated with dashed lines. (**H**) Quantification of placenta morphology from H&E sections. Shown are data points and means, n=6 placentas per genotype. Paired t-tests were used to determine significance of differences observed in different litters. Bars: G, upper panel 1mm and lower panel 0.5mm.

Placenta development involves contributions by the mother and the developing embryos, and requires dynamic cellular process, including cell migration and invasion, as well as remodeling of cell-cell junctions and cell-cell fusion; hence, placenta development is an ideal organ to study the importance of p114RhoGEF in cell dynamics. Fetal development is regulated through placental growth during mid-gestation. At E12.5, p114^−/−^ placentas were found to be significantly weight-reduced compared to wild-type littermate controls, which correlated with the reduced weight of the embryos (Fig. 1F). At E12.5, the three main layers of the placenta are the maternal decidua basalis and the embryo-derived junctional zone and labyrinth. Placentas from p114^−/−^ embryos were paler with less blood in the labyrinth zone, suggesting impaired vascularization, and the junctional zone width was reduced and had significantly fewer number of invasions into the labyrinth (Fig. 1G,H; Supplementary Fig. 1D). Thus, p114RhoGEF is required for morphogenesis of the labyrinth zone of the placenta.

### p114RhoGEF-deficiency impairs labyrinth syncytiotrophoblast differentiation

Placenta development depends on the differentiation of trophoblast stem cells (TSCs) into various subtypes of trophoblasts to form the junctional zone and the labyrinth. Markers of initial trophoblast differentiation into spongiotrophoblasts and giant cells of the junctional zone (Acsl2 and Hand1, respectively) from trophoblast stem cells (Cdx2) were not significantly affected by loss of ARHGEF18, suggesting no defect in the initial differentiation of TSCs (Fig. 2A). In contrast, markers of the labyrinth syncytiotrophoblast cells SynT-I and SynT-II (the transcription factor Gcm1 and the fusogenic proteins SynA, and SynB) were reduced in p114RhoGEF deficient placentas (Fig. 2A). Markers of TSCs and trophoblasts other than labyrinth syncytiotrophoblasts were unaltered or increased (Supplementary Fig. 2). In situ hybridization confirmed the observations made by qPCR and revealed strong expression of p114RhoGEF in the labyrinth zone of wild type but not knockout mice (Fig. 2B). Immunofluorescence indicated expression of p114RhoGEF in the labyrinth layer in both endothelia (Isolectin B4) and the neighboring syncytiotrophoblasts (MTC4, Fig. 2C,D). Hence, the expression pattern of p114RhoGEF is compatible with a role in labyrinth trophoblasts and endothelia. The attenuation of formation of labyrinth trophoblast subtypes thus provides an explanation for the defective development of the labyrinth in the absence of p114RhoGEF. SynT-I and SynT-II cells are critical for the morphogenesis of the embryonic labyrinth sinusoids, which facilitate effective fetal-maternal nutrient/gas exchange. Intriguingly, Isolectin-B4 staining, which marks blood vessels, appeared to be at similar levels in control and knockout tissue, but the vessels appeared more loosely organized in p114RhoGEF knockout mice, suggesting that blood vessels were still present (Fig. 2D). In contrast, staining for MCT4, a marker for SynT-II cells, revealed a highly organized and compact labyrinth structure in wild type placentas, but MCT4 expression was weak and disorganized in p114^−/−^ placentas (Fig. 2D). Other endothelial markers such as laminin and CD31/PECAM revealed similar changes as Isolectin-B4: a more disordered appearance but at similar expression levels. However, markers expressed by labyrinth syncytiotrophoblasts and endothelial cells (β-catenin, Vegfr2) or syncytiotrophoblasts only (E-cadherin) appeared also downregulated (Supplementary Fig. 3); syncytiotrophoblasts, but not endothelial cells, are reduced in the labyrinth layer of p114RhoGEF deficient placentas. These results indicate that p114RhoGEF plays a role in syncytiotrophoblast differentiation and blood vessel organization in the labyrinth layer of the placenta.

**Figure 2:**
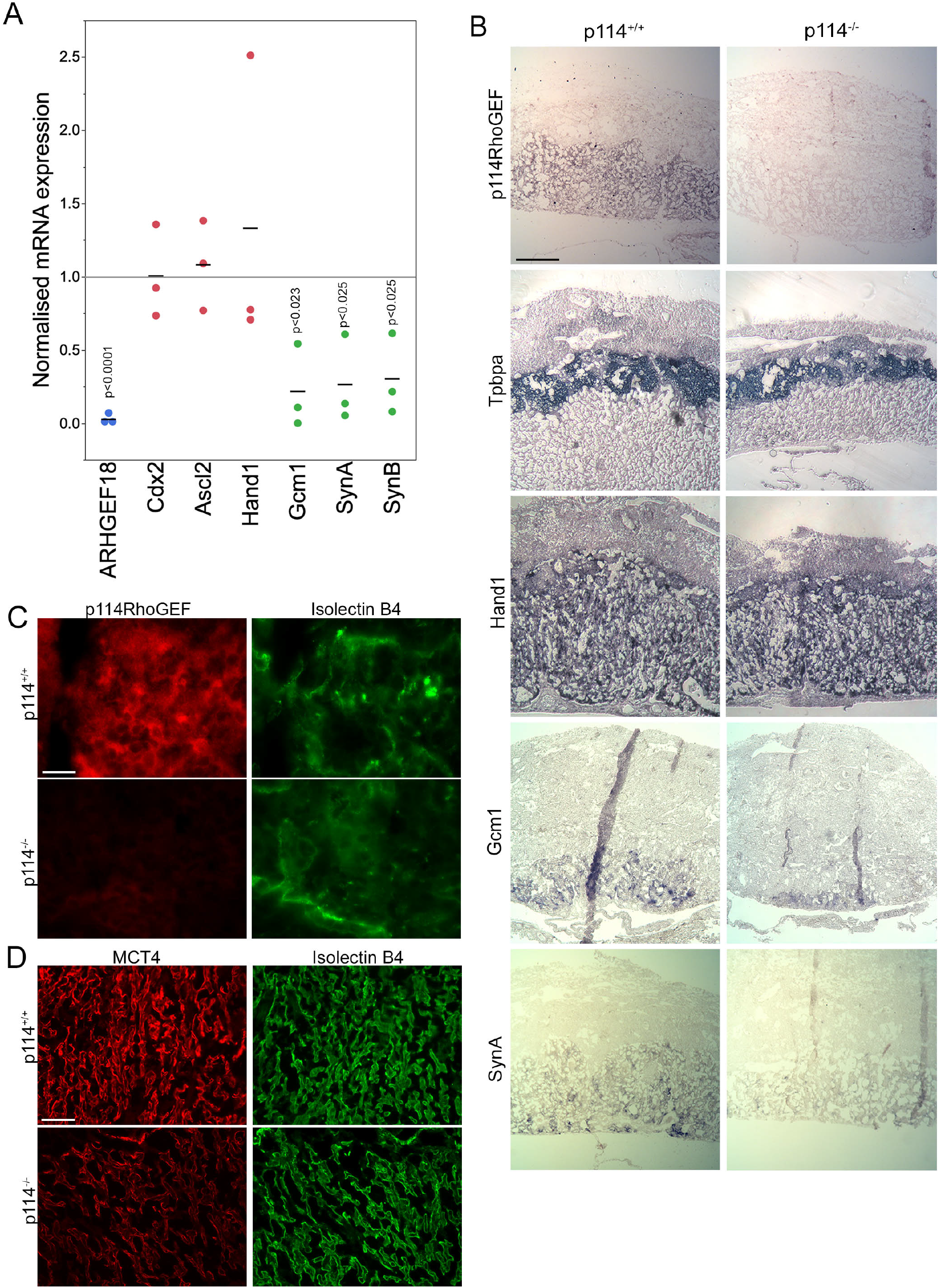
p114RhoGEF is required for labyrinth syncytiotrophoblast differentiation. (**A**) RNA expression analysis of p114RhoGEF and markers for stem cells (Cdx2), spongiotrophoblasts (Ascl2), trophoblast giant cells (Hand1) and syncytiotrophoblasts (Gcm1, SynA, and SunB;). Data were normalized to wild type controls in each litter. Shown are data points, means and p-values derived from t-tests **(B)** E12.5 control and p114^−/−^ placenta cryosections (12 μm) were analysed by in situ hybridization. p114RhoGEF was preferentially expressed in the labyrinth layer and expression was lost in knockout placentas. Of the trophoblast markers, only the syncytiotrophoblast markers Gcm1 and SynA were reduced in p114^−/−^ placentas. (**C**) Immunofluorescence analysis of p114RhoGEF expression in placenta cryosections in comparison to Isolectin B4 staining of endothelial cells. Note, p114RhoGEF staining, which is lost in knockout placentas, overlaps with but is not restricted to endothelial cells. (**D**) Staining of MCT4 and Isolectin B4 in placenta cryosections reveals a disorganized appearance of endothelia-lined blood vessels and a reduced expression of the syncytiotrophoblast marker MCT4. Bars: B, 0.5mm; C, 50μm; D, 0.2mm

### p114RhoGEF regulates endothelial actomyosin and organization of cell-cell junctions

ZO-1 recruits p114RhoGEF to cell-cell junctions in primary human microvascular endothelial cells (HDMECs) via an interaction with JACOP, a junctional adaptor protein (Tornavaca et al., 2015). Hence, the disorganization of blood vessels in the placenta labyrinth of p114RhoGEF-deficient embryos could be due to a role of p114RhoGEF in endothelial cell-cell junctions and/or to the deficiency in syncytiotrophoblasts. Therefore, we first tested the role of p114RhoGEF in cultured endothelial cells. Depletion of the Rho GEF in primary endothelial cells by RNA interference indeed induced a remodeling of the actin cytoskeleton with reduced cortical F-actin at cell-cell junctions and increased stress fibers (Fig. 3A-D). This was associated with a redistribution of active myosin, as detected by staining for double phosphorylated myosin light chain 2 (ppMLC), from cell junctions to stress fibers (Fig. 3B). While components of adherens (VE-Cadherin, p120-catenin and PECAM) and tight (ZO-1, JAM-A, and JACOP) junctions were still recruited to cell-cell contacts in p114RhoGEF depleted endothelia, junctions appeared irregular and had lost their linear structure, suggesting altered junctional tension (Fig. 3C,D). The tight junction plaque proteins ZO-1 and JACOP were affected more strongly with apparent gaps along the junctions and a strong reduction in junctional recruitment of JACOP, a binding partner of p114RhoGEF (Fig. 3B; (Tornavaca et al., 2015). These results indicate that p114RhoGEF depletion in endothelial cells *in vitro* interferes with the normal organization of cell-cell junctions and the actin cytoskeleton, and suggests a feedback mechanism in which reduced p114RhoGEF expression impacts on its own recruiting mechanism, leading to reduced junctional JACOP.

**Figure 3:**
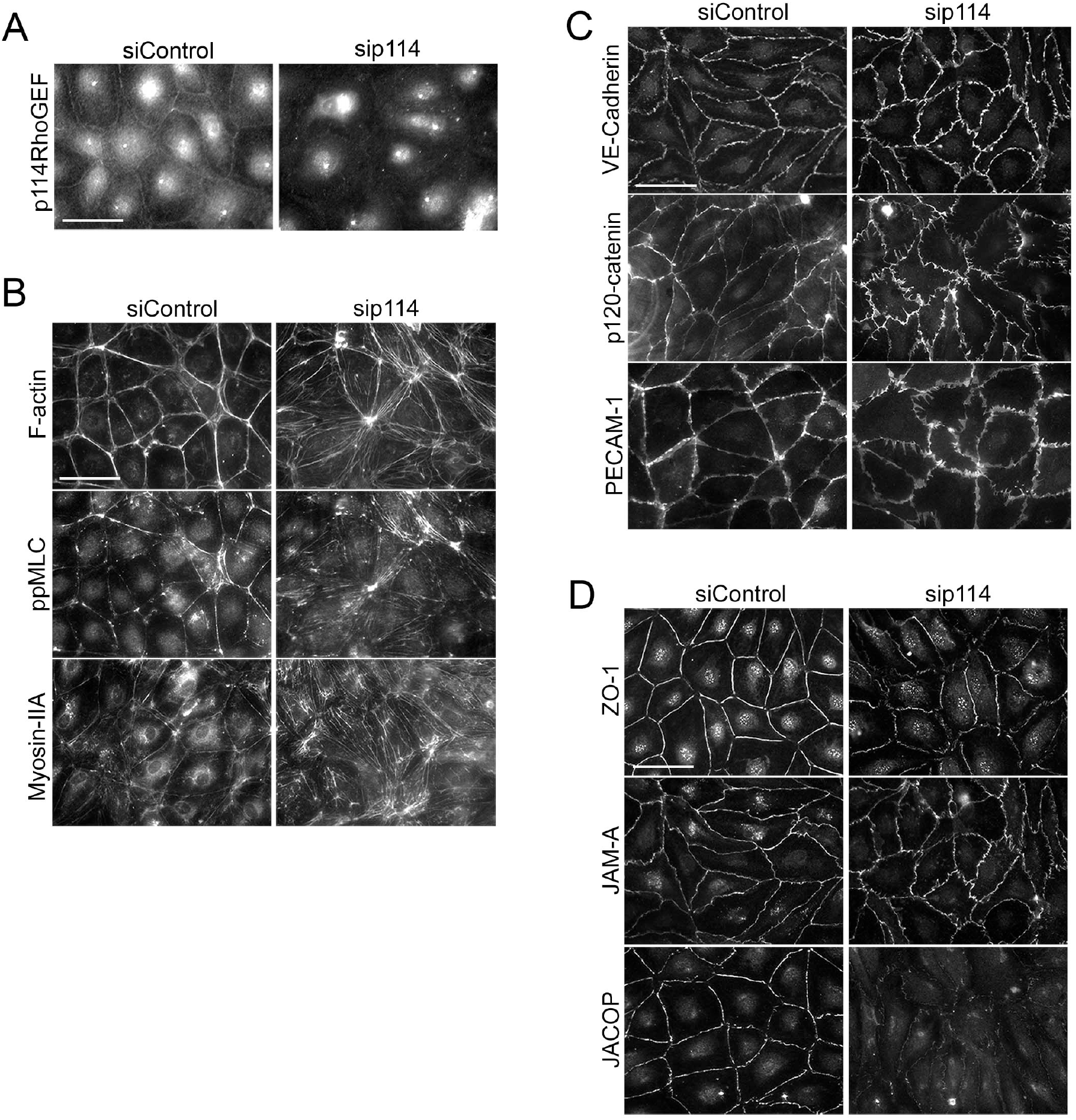
p114RhoGEF depletion in endothelial cells leads to rearrangement of the actin cytoskeleton. (**A**-**D**) HDMECs were transfected with control or p114RhoGEF-targetig siRNAs prior to analysis by immunofluorescence using antibodies against (A) 114RhoGEF; (B) double phosphorylated myosin regulatory light chain (ppMLC) and Myosin-IIA heavy chain; (C) the adherens junction proteins VE-Cadherin, p120-catenin and PECAM-1; and (D) the tight junction proteins ZO-1, JAM-A and JACOP. Fluorescently labelled phalloidin was used to label F-actin. Note, depletion of p114RhoGEF leads to reduced JACOP expression. Bars: 40μm.

### p114RhoGEF stimulates AKAP12 expression in endothelial cells and placentas but endothelial GEF is not essential for development

PKA signaling stabilizes epithelial and endothelial barriers (Matter and Balda, 2003). Given the link of p114RhoGEF to Clarkson disease (Xie et al., 2013) and the fraction of embryos with hemorrhages we observed, we asked whether p114RhoGEF cooperates with signaling proteins required for endothelial barrier stability. Several factors contribute to endothelial barrier stability with AKAP12 having been suggested to be involved in PKA-mediated barrier formation (Radeva et al., 2014). AKAP12 is a tumor suppressor gene and A kinase anchoring protein family member that regulates cytoskeletal and cell-cycle signaling pathways (Akakura et al., 2008; Radeva et al., 2014). Thus, we tested whether p114RhoGEF depletion affects AKAP12 expression. AKAP12 was downregulated by p114RhoGEF depletion as determined by RT-qPCR (Fig. 4A). By immunoblotting, the junction-associated coiled-coil protein (JACOP) was strongly reduced by p114RhoGEF depletion and more weakly upon AKAP12 knockdown (Fig. 4B). Unfortunately, we could not obtain reliable immunoblots for AKAP12 despite testing multiple antibodies, suggesting that the large size of the protein may be the underlying reason. Immunofluorescence experiments demonstrated AKAP12 localization to cell-cell junctions in control HDMECs; this signal was lost upon AKAP12 depletion with siRNAs, confirming staining specificity (Fig. 4C). As expected, the AKAP12 staining was also strongly reduced in p114RhoGEF depleted HDMECs (Fig. 4C). While the junctional staining overlapped with p114RhoGEF, AKAP12 staining was not restricted to tight junctions. Strikingly, depletion of AKAP12 by RNA interference in HDMEC led to a similar reorganization of the actomyosin cytoskeleton as p114RhoGEF depletion: F-actin was redistributed from cell-cell contacts to stress fibers and the tension sensitive protein vinculin moved from cell junctions to focal adhesions (Fig. 4D). Similarly, JACOP localization and expression was disrupted, and β-catenin staining was less linear at the junctions although not to the extent as in p114RhoGEF-depleted cells (Fig. 4C). Thus, AKAP12 is a key target and effector of p114RhoGEF signaling in endothelial junction assembly and regulation of the actin cytoskeleton. JACOP is the protein responsible for p114RhoGEF recruitment to endothelial tight junctions (Tornavaca et al., 2015). Hence, AKAP12 provides a feedback signal to p114RhoGEF by regulating expression of the component mediating junctional recruitment of the Rho GEF.

**Figure 4:**
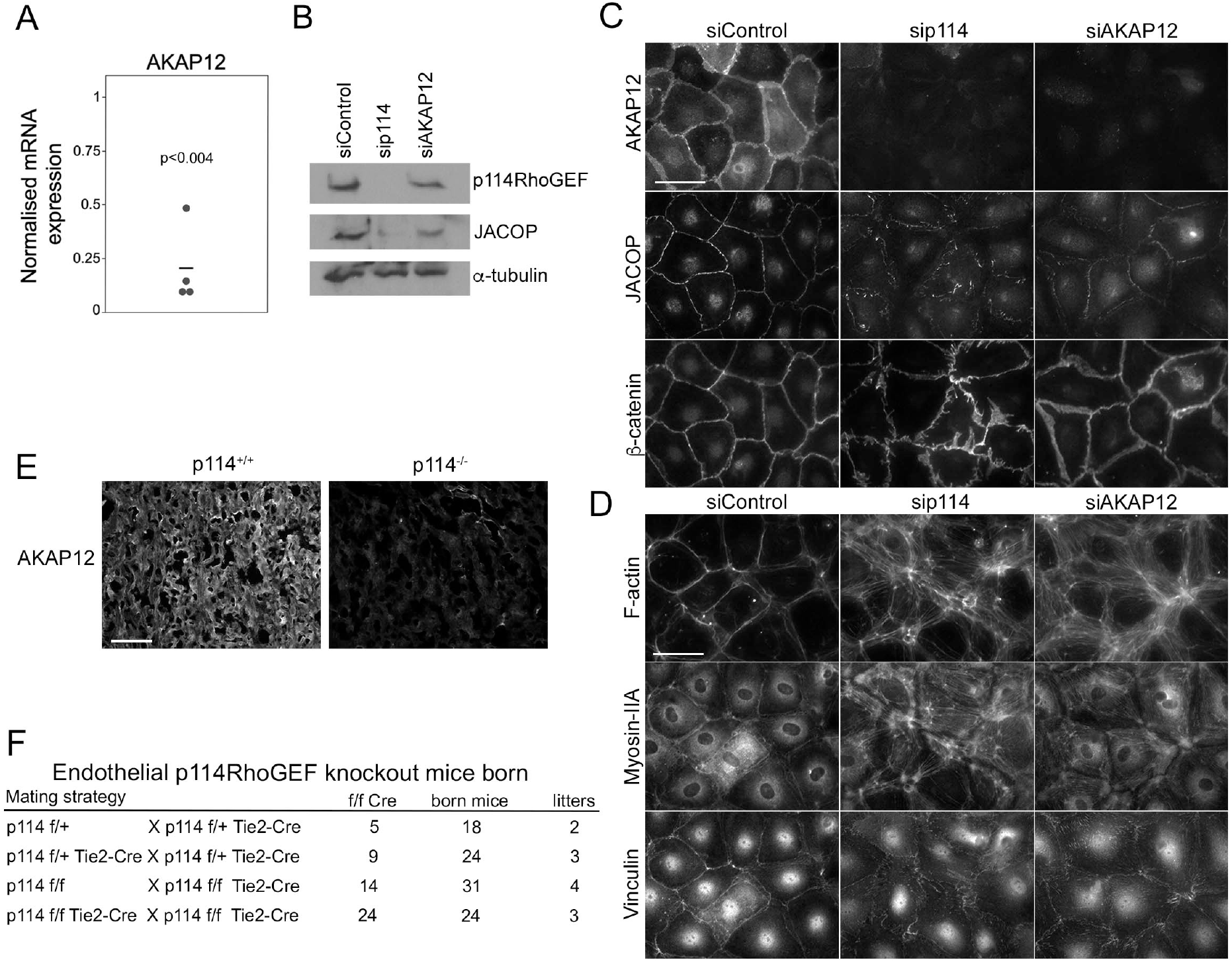
p114RhoGEF depletion suppresses AKAP12 expression in endothelial cells and placentas but endothelial specific deletion does not affect embryonic viability. (**A**) p114RhoGEF depletion reduces expression of AKAP12 mRNA as assessed by RT-qPCR. Data were normalized to wild type controls in each litter. Shown are data points, means and p-values derived from t-tests. (**B**-**D**) HDMECs were transfected with siRNAs as indicated and then processed for (B) immunoblotting or (C,D) immunofluorescence for the indicated markers. Note, AKAP12 depletion induces a strong reorganization of the actin cytoskeleton and reduced JACOP expression reminiscent of the effect of p114RhoGEF depletion. (**E**) AKAP12 staining in cryosections of placentas from wild type and p114RhoGEF deficient mice. Note the strong repression of AKAP12 expression by p114RhoGEF depletion *in vitro* and *in vivo*. (**F**) Analysis of litter genotypes derived from inter-crossing mice harboring the conditional p114RhoGEF allele (p114^f/f^) and Tie2-Cre for endothelial specific knockouts. Expected numbers of p114^f/f^ Tie2-Cre mice were born indicating that endothelial p114RhoGEF deficiency did not impede embryonic development. Bars: C and D, 40μm; E, 0.2mm.

As PKA signaling also plays an important role in the initiation of trophoblast cell fusion during placenta formation (Gerbaud and Pidoux, 2015), we next stained placenta sections for AKAP12 to determine whether downregulation of AKAP12 protein also occurred in vivo in p114RhoGEF deficient placentas. AKAP12 was found to be widely expressed in the labyrinth zone of the placenta and was strongly reduced in the absence of p114RhoGEF (Fig. 4E). Thus, AKAP12 expression is p114RhoGEF-dependent *in vitro* and *in vivo*.

The observed *in vitro* effects of p114RhoGEF depletion suggest that p114RhoGEF deficiency in mice may lead to vessel malfunction in the placenta and the embryo itself due to endothelial defects, causing the observed abdominal hemorrhages in a fraction of the knockout embryos. However, the embryonic blood vessel integrity defects were modest, arguing against a general failure of endothelial cells. To test if endothelial p114RhoGEF is required for embryonic development directly, we generated endothelial-specific p114RhoGEF knockout mice using a conditional p114RhoGEF/ARHGEF18 allele (p114^flox^) and Tie2-Cre mice. Analysis of litters born from such mice revealed the expected numbers of mice carrying the conditional allele and Tie2-Cre (Fig. 4F, Supplementary Fig. 4). Hence, endothelial specific knockout of p114RhoGEF does not lead to embryonic death.

### p114RhoGEF determines trophoblast actomyosin organization and drives migration and differentiation

As endothelial knockout of p114RhoGEF did not affect viability but the global knockout strongly affected the differentiation of labyrinth syncytiotrophoblasts during placenta development (Fig. 2), we next asked whether the RhoA activator functions in trophoblast differentiation. To study how p114RhoGEF-deficiency impacts on the regulation of trophoblast stem cell (TSC) migration and differentiation, we used the murine TSC line TS-Rs26 (Tanaka et al., 1998). Depletion of p114RhoGEF with mouse specific siRNAs induced stress fibers and redistribution of vinculin, similar to what we had observed in endothelial cells (Fig. 5A,B; Supplementary Fig. 5A). Adherens junctions were only weakly affected (Fig. 5B, Supplementary Fig. 5B). Placental development requires TCS migration; thus, we performed cell migration scratch assays on confluent monolayers. p114RhoGEF depletion by RNA interference resulted in significant reduction in wound closure by 48h (Fig. 5C), indicating a reduced migratory capacity. This is in agreement with a previous study in which we demonstrated that p114RhoGEF depletion reduces cell migration of corneal epithelial cells and invasion of tumor cell (Terry et al., 2012). Thus, p114RhoGEF regulates TSC migration providing a possible explanation for the reduced trophoblast invasions observed in knockout placentas.

**Figure 5:**
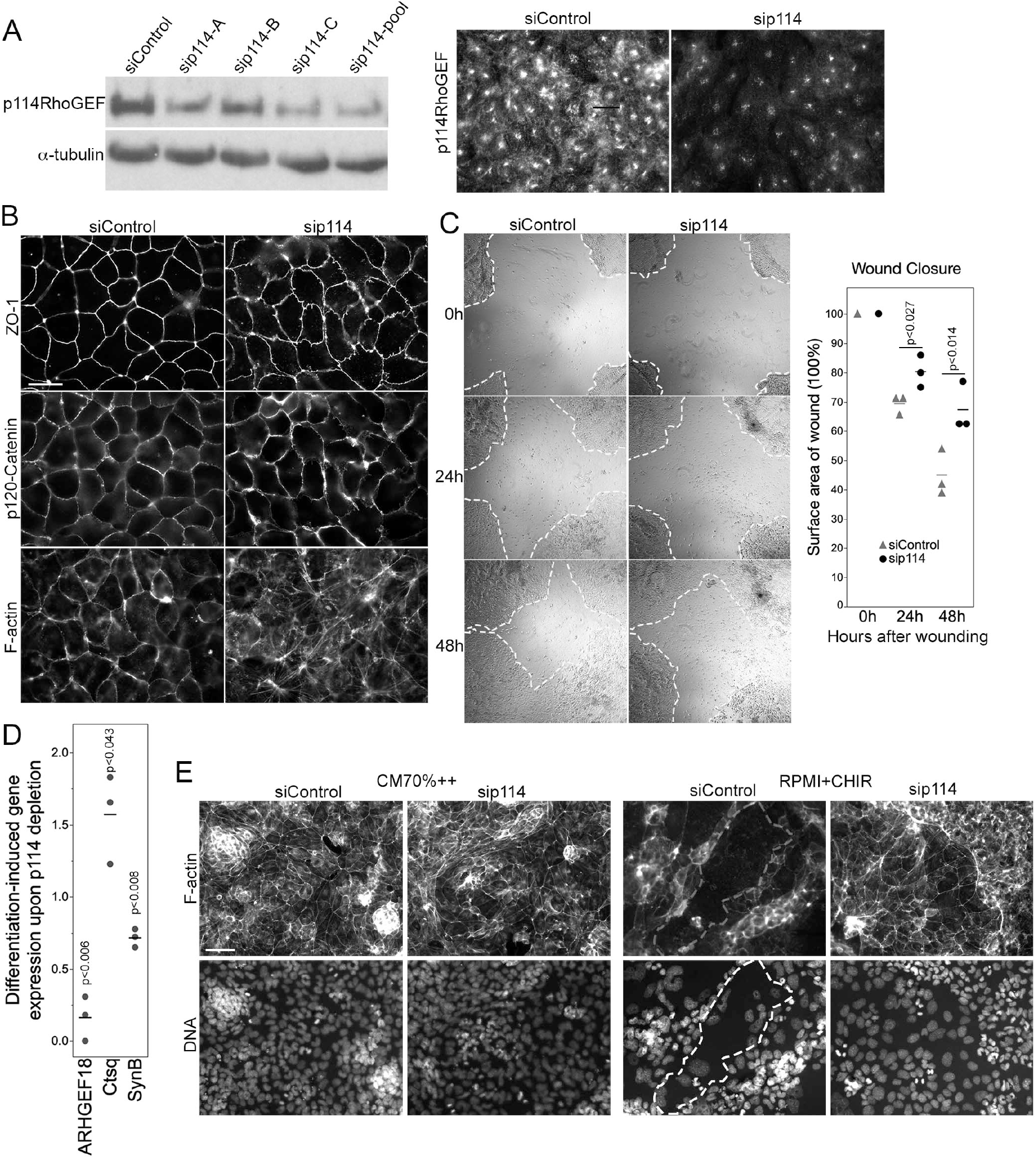
p114RhoGEF regulates trophoblast differentiation. (**A**) Depletion of p114RhoGEF in siRNA-transfected TS-Rs26 cells was analysed by immunoblotting and immunofluorescence. (**B**) Analysis of the tight junction protein ZO-1, the adherens junction protein p120-catenin, and F-actin in control and p114RhoGEF-targeting siRNA transfected TS-Rs26 cells by fluorescence microscopy. (**C**) Cell migration analysed by scratch wound closure after 0, 24 and 48h in TS-Rs26 cells transfected with siRNAs as indicated. Percentage of wound closure was calculated at each time point and plotted from three independent experiments. Shown are data points, means and p-values derived from t-tests. (**D**) Expression of trophoblast lineage markers in cells kept in standard maintenance medium (CM70%++) or upon induction of differentiation with RPMI containing the GSK3β-inhibitor CHIR was analyzed by RT-qPCR. Shown are data points of three independent experiments normalized to respective control values, means and p-values derived from t-tests. Note, depletion of p114RhoGEF attenuates induction of the syncytiotrophoblast marker SynB. (**E**) Cells cultures as in panel **D** were analyzed by fluorescence microscopy upon labelling F-actin and DNA. An area occupied by a large fused cell is demarked with a dashed line. Note, fusion was inhibited by p114RhoGEF depletion. Bars: A and B, 40μm; C, 1mm; E, 100μm.

We next asked whether p114RhoGEF-depleted TSCs can differentiate into labyrinth-specific trophoblast lineages. Induction of syncytiotrophoblast II cells (SynT-II) was achieved by GSK3β inhibition with CHIR9902 (Zhu et al., 2017). As expected, induction of differentiation resulted in downregulation of the TSC marker Cdx2 in control and p114RhoGEF-depleted cells; however, induction of SynB, a marker for SynT-II cells, was attenuated by p114RhoGEF depletion (Fig. 5D). In contrast, expression of Ctsq, a sinusoidal trophoblast giant cell marker, was enhanced in the absence of p114RhoGEF. Thus, p114RhoGEF depletion reduced SynB expression in TSCs stimulated to differentiate into labyrinth syncytiotrophoblast and in placenta *in vivo*. Induction of differentiation does not only induce expression of syncytiotrophoblast markers, but also cell-cell fusion. Strikingly, GSK3β inhibition resulted in the formation of large fused, multinucleated cells in controls but not when p114RhoGEF was depleted, suggesting that p114RhoGEF may not only be required for efficient expression of SynB but also cell-cell fusion (Fig. 5E).

### p114RhoGEF regulates cAMP-induced cell-cell fusion

Trophoblast cell-cell fusion is initiated by a cAMP signaling network (Gerbaud and Pidoux, 2015). AKAP12 is upregulated during forskolin induced cell-cell fusion although it is not known if it is required for fusion (Delidaki et al., 2011). Hence, we next asked whether p114RhoGEF and AKAP12 regulate cell-cell fusion using the human trophoblast-like BeWo cell line in which stimulation of PKA with forskolin induces cell-cell fusion and enhanced human chorionic gonadotropin (hCG) secretion (Delidaki et al., 2011). In control cells, forskolin treatment for 48h induced cell-cell fusion as observed by rearrangement of cell-cell junctions and the actin cytoskeleton that started to delineate the periphery of large multinucleated fused cells (Fig. 6A; Supplementary Fig. 6). Depletion of p114RhoGEF or AKAP12 by RNAi resulted in a strong reduction of fused cells (Fig. 6A,B; Supplementary Fig. 6). No breakdown of junctions as in control cells was observed. As expected from the *in vivo* experiments, defective fusion was paralleled by a loss in Gcm1 induction further supporting defective labyrinth - syncytiotrophoblast formation (Fig. 6C,D). Thus, p114RhoGEF and its downstream effector AKAP12 are required for cAMP-induced labyrinth trophoblast gene expression and cell-cell fusion, processes essential for syncytiotrophoblast differentiation and placenta formation.

**Figure 6:**
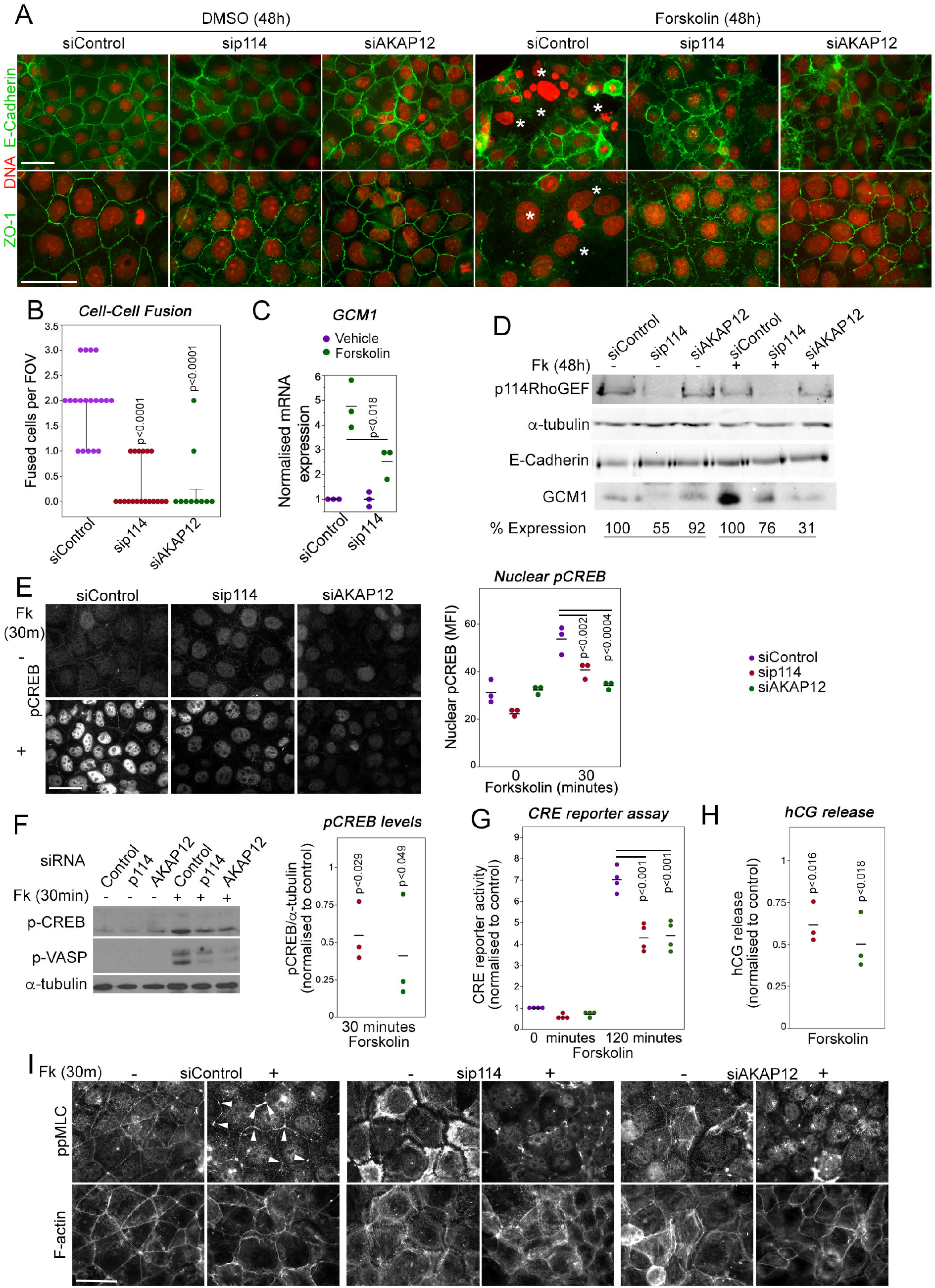
p114RhoGEF is required for forskolin-induced CREB activation, cell-cell fusion and hormonogenesis. (**A**,**B**) Control and p114RhoGEF- or AKAP12-depleted BeWo cells were treated with 100μM forskolin for 48h to induce cell-cell fusion and were then analysed by immunofluorescence microscopy after staining for DNA and E-Cadherin or ZO-1. In forskolin-treated control cells, areas of cell-cell fusion are indicated with white asterisks. Depletion of both p114RhoGEF or AKAP12 strongly reduced cell-cell fusion as supported by the quantification of fused cells per field of view (FOV) in **B** (Shown are data points, medians, interquartile ranges and p-values derived from Wilcoxon tests). (**C**) Forskolin-stimulated induction of the syncytiotrophoblast marker Gcm1 RNA is reduced in p114RhoGEF-depleted cells (shown are data points of 3 independent experiments, means and p-values derived from t-tests). (**D**) Immunoblot analysis of forskolin-induced Gcm1 protein expression. The numbers for expression levels relative to the respective control values are means derived from four experiments for p114RhoGEF and two experiments for AKAP12. (**E**) Induction of nuclear localization of pCREB by a 30-minute treatment with forskolin is inhibited by depletion of p114RhoGEF or AKAP12. The graph shows a quantification of the mean fluorescent intensity (MFI) of the nuclei (shown are data points three independent experiments, means and p-values derived from t-tests). (**F**) Immunoblots revealing induction of phosphorylation of the PKA-targets CREB (Ser-133) and VASP (Ser-239) after a 30-minute incubation with forskolin of cells transfected with either control, p114RhoGEF or AKAP12 siRNAs. The graph shows a quantification of the levels of phospho-CREB normalized to levels observed in cells transfected with control siRNAs (shown are data points of three independent experiments, means and p-values derived from t-tests). (**G**) CRE-luciferase reporter activity after a 120-minute forskolin treatment of cells transfected with siRNAs as indicated. The values were normalized to control siRNA transfected cells prior to induction for each experiment (shown are data points of four independent experiments, means and p-values derived from t-tests). (**H**) Depletion of p114RhoGEF or AKAP12 attenuates forskolin-induced secretion of hCG (shown are data point from three independent experiments, means and p-values derived from t-tests). (**I**) Depletion of p114RhoGEF or AKAP12 prevents actomyosin remodeling induced by a 30-minute forskolin treatment. Bars: 40μm.

Forskolin activation of cell-cell fusion and hormonogenesis involves PKA/CREB activation to drive expression of genes required for syncytiotrophoblast differentiation (Delidaki et al., 2011). Forskolin treatment for 30 minutes upregulated CREB phosphorylation; depletion of p114RhoGEF or AKAP12 attenuated CREB phosphorylation and nuclear staining (Fig. 6E,F). In agreement, depletion of either protein also inhibited CRE-mediated transcription in a luciferase reporter gene assay (Fig. 6G), as well as induction of human chorionic gonadotropin (hCG) release at 24 hours (Fig. 6H). Thus, p114RhoGEF and its downstream partner AKAP12 are required for efficient PKA/CREB pathway activation to induce cell-cell fusion as well as expression of PKA/CREB-dependent target genes required for syncytiotrophoblast differentiation.

Increased mechanical tension and cytoskeletal remodeling are thought to be required for cell-cell fusion (Cong et al., 2019; Kim et al., 2015). Therefore, we asked whether p114RhoGEF is required for forskolin-induced actomyosin remodeling. 30 minutes of forskolin stimulation induced preferential accumulation of activated myosin (ppMLC) at tricellular contacts (Fig. 6I). p114RhoGEF depletion led to loss of junctional actomyosin and induction of stress fibers, and forskolin stimulation was no longer able to induce enrichment of active myosin at tricellular contacts, a phenotype that was replicated by AKAP12 depletion. Similarly, depletion of either p114RhoGEF or AKAP12, reduced forskolin-induced phosphorylation of the PKA substrate VASP, a protein that regulates cytoskeletal remodeling (Fig. 6F). Thus, p114RhoGEF and AKAP12 are required for forskolin-induced actomyosin remodeling, in agreement with p114RhoGEF’s role in activation of junctional myosin contractility and the reduced phosphorylation of VASP, a regulator of actin polymerization, in cells depleted of p114RhoGEF or AKAP12.

## DISCUSSION

The RhoA activator p114RhoGEF is essential for mammalian development. A main role of p114RhoGEF during development is to drive differentiation of syncytiotrophoblasts, a cell type essential for placenta development. p114RhoGEF’s function is to coordinate actomyosin remodeling and AKAP12/PKA/CREB-driven expression of syncytiotrophoblast-specific genes to enable cell-cell fusion and syncytiotrophoblast differentiation.

Embryos deficient in p114RhoGEF exhibited strong defects in placenta morphogenesis due to defective labyrinth formation. Placenta defects are a common reason for embryonic lethality in mouse mutants (Perez-Garcia et al., 2018). Nevertheless, p114RhoGEF deficient embryos exhibited a number of phenotypes, such as hemorrhages, that are compatible with p114RhoGEF’s known role in junction formation and stability established in cell culture systems. Such phenotypes may also contribute to the observed embryonic lethality. However, mice with an endothelial specific knockout of p114RhoGEF are viable, indicating that neither embryonic not extraembryonic endothelia require the GEF to support embryonic development and placenta morphogenesis. It will be important to study the role of p114RhoGEF in junction and tissue maintenance in adult animals in response to tissue stress, which may contribute to the understanding of how p114RhoGEF malfunction leads to human disease (Arno et al., 2017; Xie et al., 2013).

Apart of the placenta, we did not observe striking morphological defects in p114RhoGEF−/− embryos. This is similar to mutant fish in which a p114RhoGEF null allele resulted in viable fish that developed an eye defect due a failure not in polarization but in maintaining cell polarity (Herder et al., 2013). Hence, p114RhoGEF is required for placenta morphogenesis but not morphogenetic processes in the embryo itself. Placenta disruption by p114RhoGEF deficiency occurred in response to defective trophoblast differentiation. In trophoblast cultures, p114RhoGEF depletion reduced cAMP-stimulated PKA/CREB activation, actin cytoskeleton rearrangements required for cell-cell fusion, and expression of genes, such as Gcm1 SynA/B, required for syncytiotrophoblast differentiation; all processes required for placenta morphogenesis. Gcm1 is a transcription factor regulating syncytiotrophoblast differentiation, and the Syn genes encode the syncytins that mediate cell-cell fusion. Our data thus indicate that p114RhoGEF coordinates AKAP12/PKA/CREP signaling and myosin activation to coordinate cytoskeletal remodeling and gene expression during cell-cell fusion and syncytiotrophoblast differentiation.

p114RhoGEF regulates junction formation by driving junctional myosin activation in cells in culture (Terry et al., 2011). Here, we identify p114RhoGEF as a regulator of transcription and F-actin formation by modulating PKA signaling and, thereby, CREB and VASP activation. *In vitro* and *in vivo* loss of function approaches indicate that the RhoA exchange factor regulates expression of AKAP12 and PKA/CREB target genes in trophoblasts that are required for syncytiotrophoblast fusion (e.g., Gcm1, SynB, hCG) by attenuating phosphorylation and, thereby, activation of CREB. The spatial regulator of PKA signaling AKAP12 is downregulated by p114RhoGEF depletion in *in vitro* models and in the placenta *in vivo*. While AKAPs are thought to act as scaffolding proteins that determine the spatial organization of PKA signaling, how p114RhoGEF regulates expression of AKAP12 is not clear yet.

Active p114RhoGEF forms complexes with Rock II and myosin-IIA, which is important for its role in junction formation and cell migration (Terry et al., 2010; Terry et al., 2012). Interestingly, both Rock II and myosin-IIA are important for placenta development. Knockout of myosin-IIA was reported to impact on an earlier stage of placenta development than p114RhoGEF deficiency, suggesting that different mechanisms of actomyosin activation are involved at different stages of trophoblast linage differentiation (Crish et al., 2013). Rock II is highly expressed in the labyrinth layer and its deletion leads to disruption of the architecture of the labyrinth layer (Thumkeo et al., 2003). While overall F-actin architecture in the labyrinth layer and stress fiber formation in trophoblasts in culture were not affected by Rock II deficiency, myosin activity and cell-cell junctions in trophoblasts were not studied in the Rock II knockout animals. Hence, it is not clear how Rock II contributes to actomyosin activity in trophoblasts; however, a role in gene expression was suggested (Thumkeo et al., 2003). As our results indicate that p114RhoGEF, which complexes with Rock II, regulates trophoblast gene expression, the two proteins may cooperate in at least in some functions during trophoblast differentiation. The GTPase cycle not only requires GEFs but also GAPs. DLC1, a GAP for RhoA and Cdc42, is highly expressed in trophoblasts of the labyrinth, and DLC1 knockout was reported to lead to defects in labyrinth organization; however, the underlying reasons have not been analysed (Durkin et al., 2005). It is possible, that DLC1 may be a signaling partner of p114RhoGEF in some processes as DLC1 was reported to stabilize cell-cell junctions in cultured tumor cells (Tripathi et al., 2012). Another signaling partner of p114RhoGEF, LKB1, has also been linked to placenta development, at least in part due upregulation of VEGF secretion (Ylikorkala et al., 2001). In contrast to myosin IIA and Rock II, LKB1 functions upstream of p114RhoGEF by binding and activating the GEF in a kinase activity-independent manner, which is important for junction formation and morphogenesis of cultured epithelial cells (Xu et al., 2013). However, whether LKB1 plays a role in trophoblasts and cell-cell fusion is not known.

Other cell-cell junction proteins have previously been linked to placenta development. Depletion of the cell-cell adhesion genes connexin-31 or connexin-43 results in placental abnormalities and developmental defects (Kibschull et al., 2005),(Dunk et al., 2012). Connexin-43 has been shown to regulate cell-cell fusion and to form a complex with ZO-1 in trophoblasts (Pidoux et al., 2014). ZO-1 is required for junctional recruitment of p114RhoGEF and, in primary endothelial cells, ZO-1 depletion reduces cell-cell tension due to loss of junctional p114RhoGEF (Terry et al., 2011; Tornavaca et al., 2015). Although p114RhoGEF depletion in trophoblast models did not prevent junctional recruitment of ZO-1, the morphological appearance of the junctional staining of ZO-1 indicated reduced cell-cell tension. This was further supported by the observed striking remodeling of the actomyosin cytoskeleton in p114RhoGEF-deficient trophoblasts with a strong reduction in junction-associated active myosin and an induction of stress fibers. Thus, it is possible that the placenta defect of Connexin-43 depletion is mediated by reduced junctional recruitment of p114RhoGEF.

Strikingly, p114RhoGEF was not only required for PKA-stimulated cell-cell fusion but also for the rapid induction of junctional foci of active myosin. Cytoskeletal remodeling and the thereby generated mechanical tension are thought to be key drivers of cell-cell fusion by promoting the close proximity of the neighboring plasma membranes required for fusion to occur (Cong et al., 2019; Kim et al., 2015). p114RhoGEF is required for tight junction assembly when monolayers are under high mechanical tension (Haas et al., 2020). As p114RhoGEF is required for PKA-stimulated remodeling of the actin cytoskeleton along cell-cell junctions prior to fusion, p114RhoGEF-deficient trophoblasts may not be able to generate and maintain the necessary close membrane-membrane contacts required for membrane fusion. Hence, p114RhoGEF drives placenta morphogenesis and syncytiotrophoblast differentiation by modulating PKA-stimulated expression of genes as well as by remodeling the actomyosin cytoskeleton to enable cell-cell fusion.

## ACKNOWLEDGEMENTS

This work was supported by the BBSRC (BB/N001133/1 and BB/N014855/1). We would like to thank to Professors James Cross for providing plasmids for in situ hybridization, Marcus Fruttiger for technical advice, Christiana Ruhrberg for the Tie2-Cre mice, and Myriam Hemberger for the murine TSC line TS-Rs26 and critical reading of the manuscript.

## AUTHOR CONTRIBUTIONS

RB, KM and MSB performed experiments. AAF assessed p114RhoGEF expression in the endothelial specific knockout mice. DKG assessed β-HCG production. MSB and KM designed the project. All authors read and contributed to the final version of the manuscript.

## COMPETING INTERESTS

The authors declare no competing interests.

## MATERIALS AND METHODS

### Mouse lines

ARHGEF18/p114RhoGEF knockout mice containing a gene trap insertion ablating expression were purchased from (EMMA - https://www.taconic.com/knockout-mouse/arhgef18-trapped) and animals carrying the conditional knockout first (promoter driven) ARHGEF18^tm1a(KOMP)^ allele were obtained from the Knockout Mouse Project (KOMP; https://www.komp.org/geneinfo.php?geneid=23667). ARHGEF18/p114RhoGEF knockout mice containing a gene trap insertion ablating expression were crossed into a C57BL/6N genetic background for more than 6 generations. Animals carrying the conditional knockout first (promoter driven) ARHGEF18^tm1a(KOMP)^ allele were also crossed into a C57BL/6N genetic background. The lacZ/neo cassette was removed by crossing with mice carrying the Flippase gene. To generate endothelial-specific knockouts, animals carrying the conditional allele were crossed with mice harboring the Tie2-Cre driver (kindly provided by Professor Christiana Ruhrberg) (Erskine et al., 2017; Kisanuki et al., 2001). Animals were housed in individually ventilated cages and facilities were regularly monitored for health status. Use of all animals was in accordance with UK Home Office regulations, the UK Animals (Scientific Procedures) Act of 1986 and was approved by the Institute’s Animal Welfare and Ethical Review Body. The number of mice per experimental group was kept to the minimum to reach statistical significance and ensure reproducibility in accordance with NC3R recommendations. Timed pregnancies were monitored by counting the day of the vaginal plug as E0.5 and pregnant females were sacrificed and dissected from E10.5 through E15.5. The pregnant mice were first euthanized and their uteri were removed by cutting at the cervix. Placentas and embryos were collected and imaged for phenotyping. Tail samples were collected for genotyping by PCR using genomic DNA and primer 5’-ATCCAGTAACTACCATACCCACCC-3’ together with primer 5’-GGCTTAGACGAACAGGAGTTCCAAG - 3’ for the wild type allele and with primer 5’-TATTCAGCTGTTCCATCTGTTCCTGACC - 3’ for the mutant allele. The floxed allele was detected with 5’-ATTTTTGTCTGCATGTATGTCTGTGC-3’ and 5’-GAGATGGCGCAACGCAATTAATG-3’, and Tie2-Cre with 5’-GCCTGCATTACCGGTCGATGC-3’ and 5’-CAGGGTGTTATAAGCAATCCCC-3’. Placentas and embryos were imaged with a Nikon SMZ1500 stereomicroscope equipped with a Plan Apo 0.5x lens and a DS-Fi2 camera controlled with a DS-L3 unit.

### Cell lines

Human dermal microvascular endothelial cells were purchased form Promocell (HDMEC-c adult C-12212). The wild-type mouse TS-Rs26 trophoblast stem cell (TSC) line was kindly provided by Professor Myriam Hemberger (Latos et al., 2015; Murray et al., 2016). BeWo cells were described previously (Delidaki et al., 2011). Fresh batches of cells from a contamination-free stock that had been tested for mycoplasma were used to replace fresh cultures every 6 to 8 weeks. Cells were then weekly stained with Hoechst dye to reveal nuclei and DNA of contaminants such as mycoplasma.

### Histology analysis, immunofluorescence and immunoblotting

Placentas were fixed in 4% PFA, frozen in OCT-embedding compound and cryo-sectioned. Consecutive 12μm cryosections were produced and alternately mounted on different slides. A series of sections per block was processed for hematoxylin and eosin (H&E) staining. For immunofluorescence, cryosections were permeabilized with 0.5% Triton X-100 and 0.1% Saponin in PBS and then blocked with 1% donkey serum in 0.1% TritonX-100 and 0.02% Saponin in PBS. Primary antibodies were incubated overnight at 4°C in blocking solution. Secondary antibodies were then also incubated in blocking solution, followed by washing with PBS and mounting with Prolong Gold antifade reagent (Life technologies). Cells were fixed either with methanol at −20C° for 5 minutes, 95% ethanol for 10 minutes at −20C°, or with 3% PFA for 20 minutes at room temperature followed by 0.3% Triton X-100 permeabilization for 5 minutes (Balda et al., 1996). The cells were then washed with PBS and incubated in blocking buffer (PBS containing 0.5% BSA) for 15 minutes prior to labelling with primary antibodies in the same buffer (Tornavaca et al., 2015). After an overnight incubation at 4°C, cells were washed three times with blocking buffer and then incubated with the secondary antibodies in the same buffer for at least 2 h. After three washes with PBS, the coverslips were mounted with ProLong Gold mounting medium (Invitrogen) and stored at 4°C (Elbediwy et al., 2012). Images were taken with an epifluorescence microscope (Zeiss Axioscope) using a 40×/1.4 NA objective lens and a Hamamatsu Photonics camera C4742-95 camera using and simple PCI software (Hamamatsu Photonics). Brightness and contrast were adjusted with Photoshop CS4 and CC (Adobe). Fluorescence intensity and placenta parameters were quantified with ImageJ/Fuji software. For immunoblotting, cells were washed twice with PBS, lysed in SDS-PAGE sample buffer, and denatured at 70°C for 10 min (Sourisseau et al., 2006). The samples were then homogenized with a 1-ml syringe and a 25G needle, proteins were separated by SDS-PAGE and transferred onto PVDF membranes (Steed et al., 2014). For immunoblotting of E12.5 placentas and embryos, whole tissue samples were homogenized in 6M Urea, protein content quantified, diluted in 3XSDS-PAGE sample buffer and separated in mini-gels. The membranes were blocked with 5% defatted milk powder dissolved in TBS containing 0.1% Tween-20 and then incubated with primary antibodies overnight in the same solution or, for anti-phospho-protein antibodies, 5% BSA dissolved in TBS containing 0.1% Tween-20. After washing, the primary antibodies were detected with HRP-conjugated or fluorescent secondary antibodies using either an ECL detection system or Li-Cor ODYSSEY infrared imaging system (Steed et al., 2014).

### In situ hybridization

In situ hybridization was performed in 12-μm placenta cryosections using digoxygenin (DIG) labelled probes and an anti-DIG-alkaline phosphatase detection system. Probes for Tpbpa, Hand1, Gcm1 and SynA were kindly provided by Professor James Cross and used as previously described (Simmons et al., 2007). The p114RhoGEF probe for in situ hybridization was custom-designed to be used with the ViewRNA system, which was used according to the manufacturer’s instructions (ThermoFisher). Briefly, frozen OCT 12μm placenta sections were dehydrated through ethanol (50%-70%-100%) and then baked dry for 1 hour at 60°C. Samples were then treated with proteinase for 5 minutes, fixed in 4% formaldehyde and then incubated with p114RhoGEF viewRNA probe overnight at 40°C in a humidified chamber. The following day, samples were incubated with pre-amplification and amplification probes for 1 hour and 30 minutes, respectively, at 40°C. A final step to label the reaction with alkaline phosphatase was performed at 40°C. Samples were then subjected to a final wash before incubation with substrate overnight to allow purple precipitate to develop.

### Cell culture and RNA interference

Human dermal microvascular endothelial cells were maintained on gelatin (G1393; Sigma-Aldrich)-coated tissue-culture Petri dishes in endothelial cell growth Medium MV2 (Promocell) (Tornavaca et al., 2015). Cells were used between passage two and four. The wild-type mouse TS-Rs26 trophoblast stem cell (TSC) line was kindly provided by Professor Myriam Hemberger and cultured as described (Latos et al., 2015; Murray et al., 2016). In brief, TS-Rs26 TSC line was maintained in 70% volume mouse embryonic fibroblasts (MEF) conditioned medium (CM, collected from Mitomycin-C treated MEFs), 30% volume RPMI-1640 containing 20% fetal bovine serum, 1% non-essential amino acids, 1% sodium pyruvate and 0.1% β-mercaptoethanol, supplemented with FGF 50 ng/ml and Heparin 1 μg/ml as previously described (Murray et al., 2016). BeWo cells were maintained in 50% F12K/50% DMEM medium supplemented with 15% fetal bovine serum. Cells were passaged every 3-4 days at subconfluency and medium was changed daily. For RNA interference, siRNA stocks were diluted to 20μM and were transfected with Lipofectamine RNAiMax reagent (Invitrogen) according to manufacturer’s instructions using final siRNA concentration of 20–80 nM. Cells were collected for analysis after 72-96 hours post-transfection. The following siRNA sequences were used: human p114RhoGEF 5’ - UCAGGGCGCUUGAAAGAUA - 3’ and 5’ – GGACGCA ACUCGGACCAAU - 3’; mouse p114RhoGEF 5’ - GCAUCAUCCAGAACACAGA - 3’, 5’ – CAGAUUCUCAGAUCGGCCA - 3’ and 5’ - CACAUGAGUUUGAGGCCGA - 3’; and human AKAP12 5’ - UCUGCAGAAUCUCCGACUA - 3’ and 5’ - AGACGGAUGUAGUGUUGAA - 3’. Non-targeting control siRNAs were 5’-UGGUUUACAUGUCGACUAA-3’ and 5’-UGGUUU ACAUGUUGUGUGA-3’. All siRNAs were synthesized by Sigma-Aldrich with dTdT 3’-overhangs.

### Stem cell differentiation, cell fusion and migration

To induce TSC line cell differentiation, cells were cultured in RPMI-1640 containing 1μM CHIR99021 inhibitor for 72h. To induce BeWo cell-cell fusion, cells were treated with the adenyl cyclase activator forskolin (100μM) for the indicated times. DMSO was used to dissolve CHIR99021 and forskolin and used as solvent control. For analysis of migration, TSCs were grown to confluency and a cross-scratch wound was induced in the monolayer using a pipette tip, and debris was removed by washing once in culture medium. Cell monolayers were then imaged after 0, 24 and 48 hours to observe wound closure. Percentage wound area was calculated by manual tracking of the wound edges using Fiji/ImageJ software. In each of the repeat experiments, three scratch wounds were analyzed.

### RNA isolation and qPCR

After 72-96h protein depletion by siRNA, cells were washed once with PBS and then lysed for RNA extraction using the PureLink RNA Mini Kit (ThermoFisher). cDNA was synthesized using the QuantiTect Reverse Transcription Kit (Qiagen). Real time quantitative PCR (qPCR) was performed using PowerUp SybrGreen Master Mix (ThermoFisher) and the QuantStudio 6 Real-Time PCR System (Applied Biosystems). GAPDH was amplified as a control to normalize the data. The following primers were used: mouse SynA 5’ – CTCCAGGAGGCTAACTCTTCC - 3’ and 5’ - TCCGGGCTGAGTACATGATTC - 3’; mouse SynB 5’ - TGGGTCCTCTGTTTCGTCCTT - 3’ and 5’ - GGGAAGAGTTGGTATCACGTAGG - 3’; mouse Gcm1 5’ - AGAGATACTGAGCTGGGACATT - 3’ and 5’ - CTGTCGTCCGAGCTGTAGATG - 3’; mouse ARHGEF18 5’ - TCAGACAGAAGTGTGGTCCG - 3’ and 5’ - GGAGACTGCGAGAGCGAC - 3’ or 5’ – TTGTGCGAAGGCTGGGAG - 3’ and 5’ – GGATGATGCGTTCCACAAGC - 3’; mouse Tpbpa 5’ - ACTCCCAGGCATAGGATGAC - 3’ and 5’ - TGAAGAGCTGAACCACTGGA - 3’; mouse Hand1 5’ – CTTTAATCCTCTTCTCGCCG - 3’ and 5’ - TGAACTCAAAAAGACGGATGG - 3’; mouse Cdx2 5’ - TCTGTGTACACCACCCGGTA - 3’ and 5’ - GAAACCTGTGCGAGTGGATG - 3’; mouse Ctsq 5’ - GTGTTTCAGCATTTGATCCCAGT - 3’ and 5’ - GTCAGCAAACCCATTTAATCCCA - 3’; mouse Ehox 5’ - GGTGATGCAGACCTCATGGAT - 3’ and 5’ - GATACCAGCACTGGAATAGGC - 3’; mouse Elf5 5’ - CTGGTCACAGCAGAATTGGA - 3’ and 5’ - CTGCCTTTGAGCATCAGACA - 3’; Ascl2 5’ - AAGCACACCTTGACTGGTACG - 3’ and 5’ - AAGTGGACGTTTGCACCTTCA - 3’; mouse Tcfeb 5’ - GGAGCCAGAGCTGCTTGTTA - 3’ and 5’ - CAAGGCCTCTGTGGATTACA - 3’; mouse Mdfi 5’ - CTGGGACCTGGAGAAAACAG - 3’ and 5’ - CGCAGCTTGCACGAGTATG - 3’; mouse Prl3d1 5’ - GGTGTCAAGCCTACTCCTTTG - 3’ and 5’ - GTATTATGGAGCAGTTCAGCCAA - 3’; mouse Prl3b1 5’ - CACCAGACAACATCGGAAGAC - 3’ and 5’ - TGACAGCAGAGTATCAGGTACA - 3’; mouse Prl2c2 5’ – TCCTGGATACTGCTCCTACTACT - 3’ and 5’ - AGCCCAGACACGTTAGAATAATG - 3’; human AKAP12 5’ - GGACCCCCTTTCTGAGAGAC - 3’ and 5’ - CAGACACCACCGCGGAC - 3’; human Gcm1 5’ - TTCCCGGTCACCAACTTCTG - 3’ and 5’ - GTAAACTCCCCTGACTTTGTGTT - 3’; human GAPDH 5’ - TTGATGGCAACAATCTCCAC - 3’ and 5’ - CGTCCCGTAGACAAAATGGT- 3’.

### Reporter assays

BeWo cells were plated into 96 well dishes at 5,000 cells per well and after 24 hours transfected with control, p114RhoGEF or AKAP12 siRNAs. To measure CREB activity, a plasmid containing a cAMP response element binding site driving firefly luciferase was transfected with a control plasmid containing a CMV promotor driving renilla luciferase expression using the TransiT-X2 transfection reagent (Mirus Bio) 48 hours after the siRNA transfection. Firefly and renilla luciferase activity were measured 24 hours later using the dual luciferase assay kit (Promega).

### Human chorionic gonadotropin (hCG) secretion assay

BeWo cells were grown in 6-well plates until a confluency of about 70%. Cells were then incubated for 24 h prior to hormone collection. Control cultures received the DMSO vehicle in the same concentration as the forskolin-treated cultures. The medium was collected after 24 h of incubation. Supernatants were aspirated, centrifuged at 500g for 5 min at 4°C to remove cell debris, and stored at −80°C until analysis. The amount of secreted hCG was determined by the Department of Biochemistry (University Hospitals Coventry and Warwickshire NHS Trust) using the Elecsys^®^ Intact hCG+b electrochemiluminescence immunoassay (ECLIA) and the fully automated modular analytics E170 testing system from Roche Diagnostics (Mannheim, Germany). Results were expressed as IU/ml per 10^5^ cells.

### Statistics and reproducibility

For the quantifications shown, the provided n values refer to the numbers of embryos or placentas analyzed in the mouse experiments or, for the experiments with cell lines, the number of repeat experiments. The data shown are individual data points along with a value representing the median or mean as indicated. Statistical significance was tested with ANOVA and two-tailed t-tests, or Kruskal-Wallis and Wilcoxon nonparametric tests as indicated using either Prism or JMP Pro (V14) software.

## Notes

### Competing Interest Statement

The authors have declared no competing interest.

