## Supplemental figures for "ARHGEF18/p114RhoGEF coordinates PKA/CREB signaling and actomyosin remodeling to drive trophoblast cell-cell fusion during placenta morphogenesis"

### SUPPLEMENTARY FIGURES

A

Genotypes of born mice

| Mating strategy | +/+ | +/- | -/- |
|---|---|---|---|
| p114♀+/+ X p114♂+/- | 52 | 47 | - |
| p114♀+/- X p114♂+/+ | 33 | 35 | - |
| p114♀+/- X p114♂+/- | 12 | 30 | 0 |

B

| | +/+ | +/- | -/- |
|---|---|---|---|
| ***E10.5-11.5*** | 27 | 62 | 22 |
| Turn defect | 3.7% | 1.6% | 18.2% |
| Effusion | - | 1.6% | 13.6% |
| Yolk sac oedema | 7.4% | 6.5% | 9.1% |
| Anemia | - | 3.2% | 4.5% |
| Resorbed | 3.7% | 6.5% | 4.5% |

C

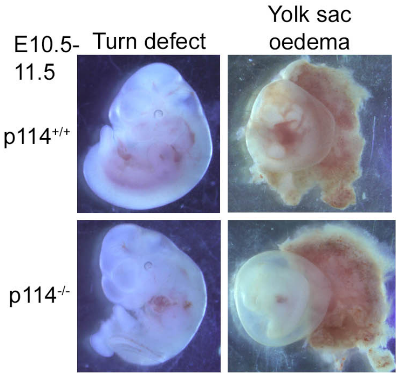


D

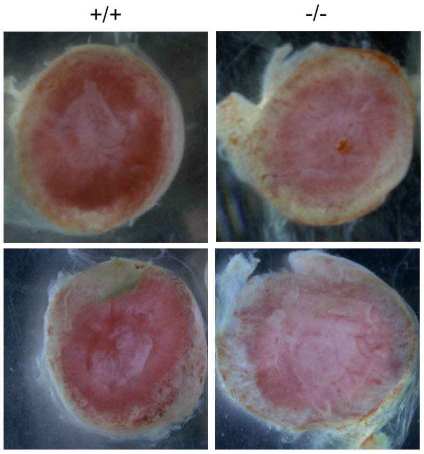


**Supplementary Figure 1: Phenotypic analysis of p114RhoGEF-deficient mice.**
(**A**) Analysis of litter genotypes derived from inter-crossing of p114$^{+/-}$ with p114$^{+/+}$ or p114$^{+/-}$ mice; no p114$^{-/-}$ mice were born. (**B,C**) Analysis of p114$^{-/-}$ phenotypes at E10.5-11.5 and representative images. (**D**) Representative images of p114$^{-/-}$ placentas at E12.5 suggesting reduced blood content.

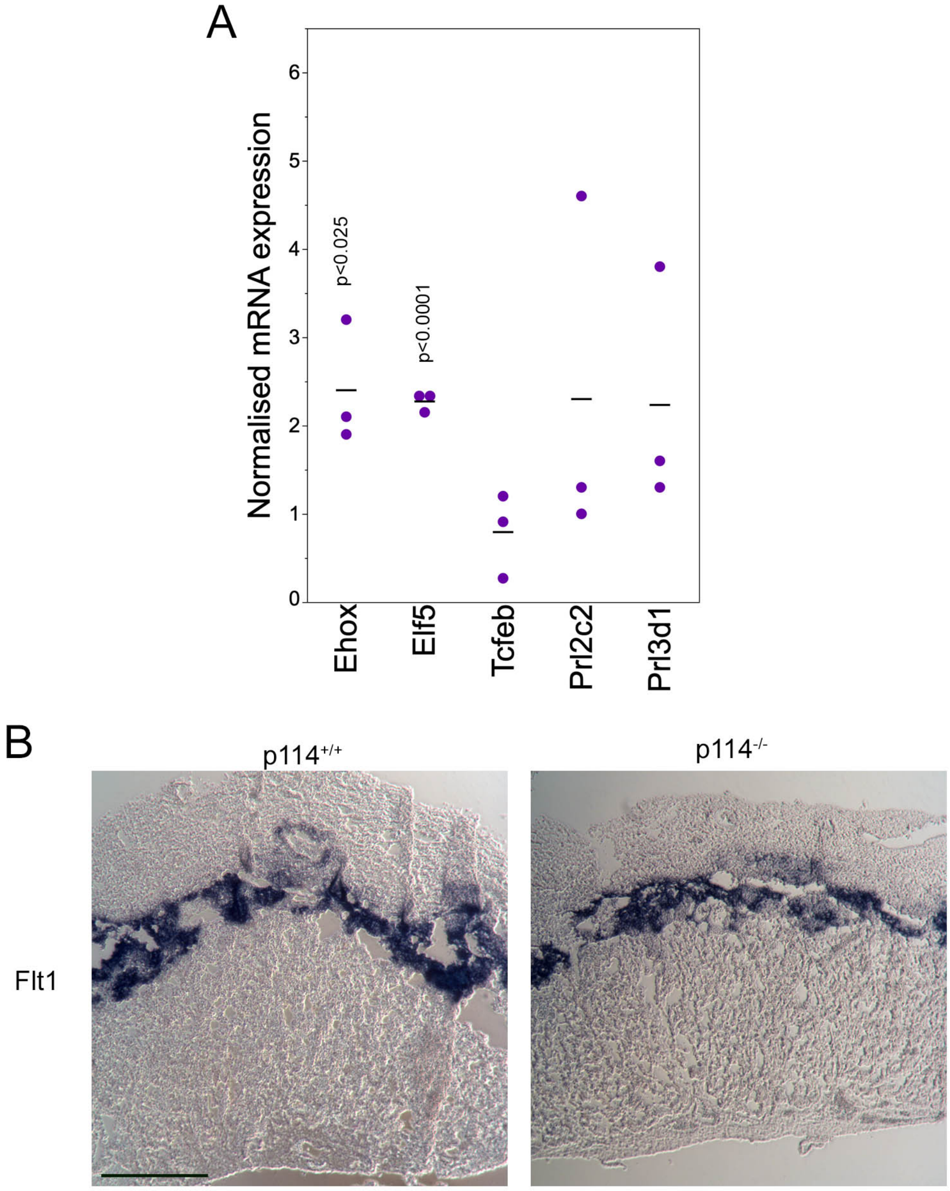


**Supplementary Figure 2: Characterization of TSC and trophoblast marker expression in p114RhoGEF-deficient mice.**

(**A**) Analysis of RNA expression by RT-qPCR of markers for TSCs and trophoblasts (shown are data points from three experiments, means and p-values derived from t-tests). (**B**) In situ hybridization revealing expression of Flt1 in the junctional zone. Bar: 0.5mm

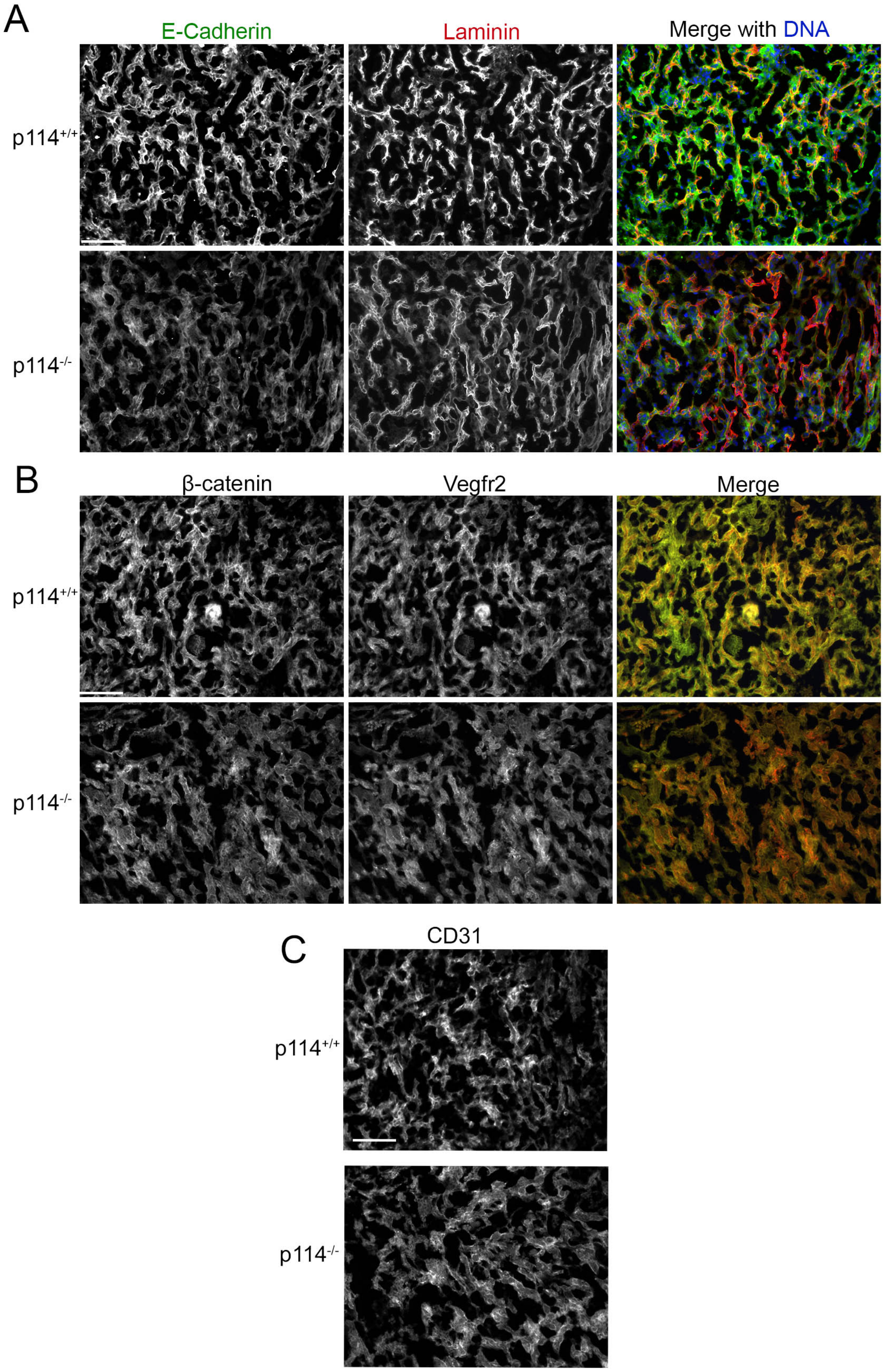


**Supplementary Figure 3: Effect of p114RhoGEF knockout on the expression of proteins of endothelia and syncytiotrophoblasts in the labyrinth layer of the placenta.**

(**A**-**C**) Expression of markers for endothelial and syncytiotrophoblasts was analyzed by immunofluorescence of cryosections. Bars: 200µm.

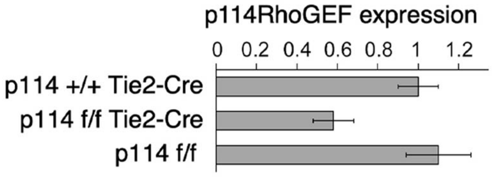


**Supplementary Figure 4: Reduced expression of p114RhoGF in aorta samples of endothelial specific p114RhoGEF knockout mice.**
Analysis of p114RhoGEF expression by RT-qPCR from total RNA isolated from aorta samples derived from control or endothelial specific knockout mice. Shown are examples of mice analysed.

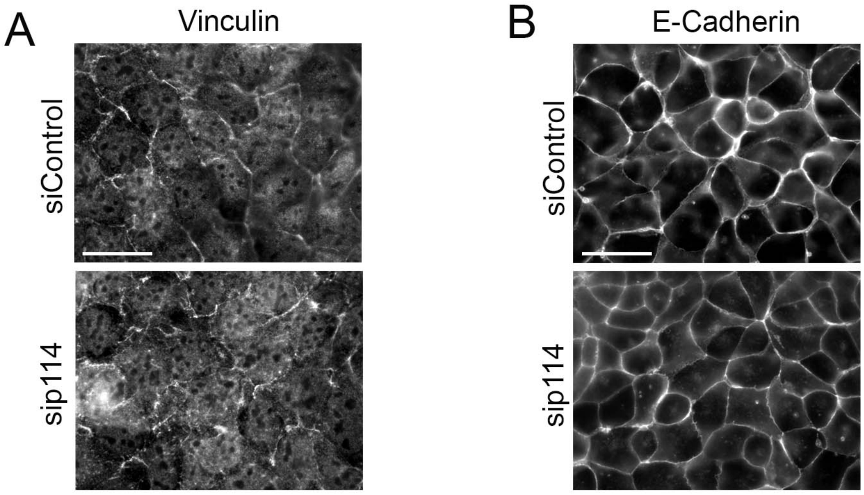


**Supplementary Figure 5: Localization of vinculin and E-cadherin in TSR-26 cells.**
Effect of p114RhoGEF depletion on vinculin (**A**) and the adherens junction protein E-cadherin (**B**) in TSR-26 cells was analyzed by immunofluorescence microscopy. Bars: 40μm.

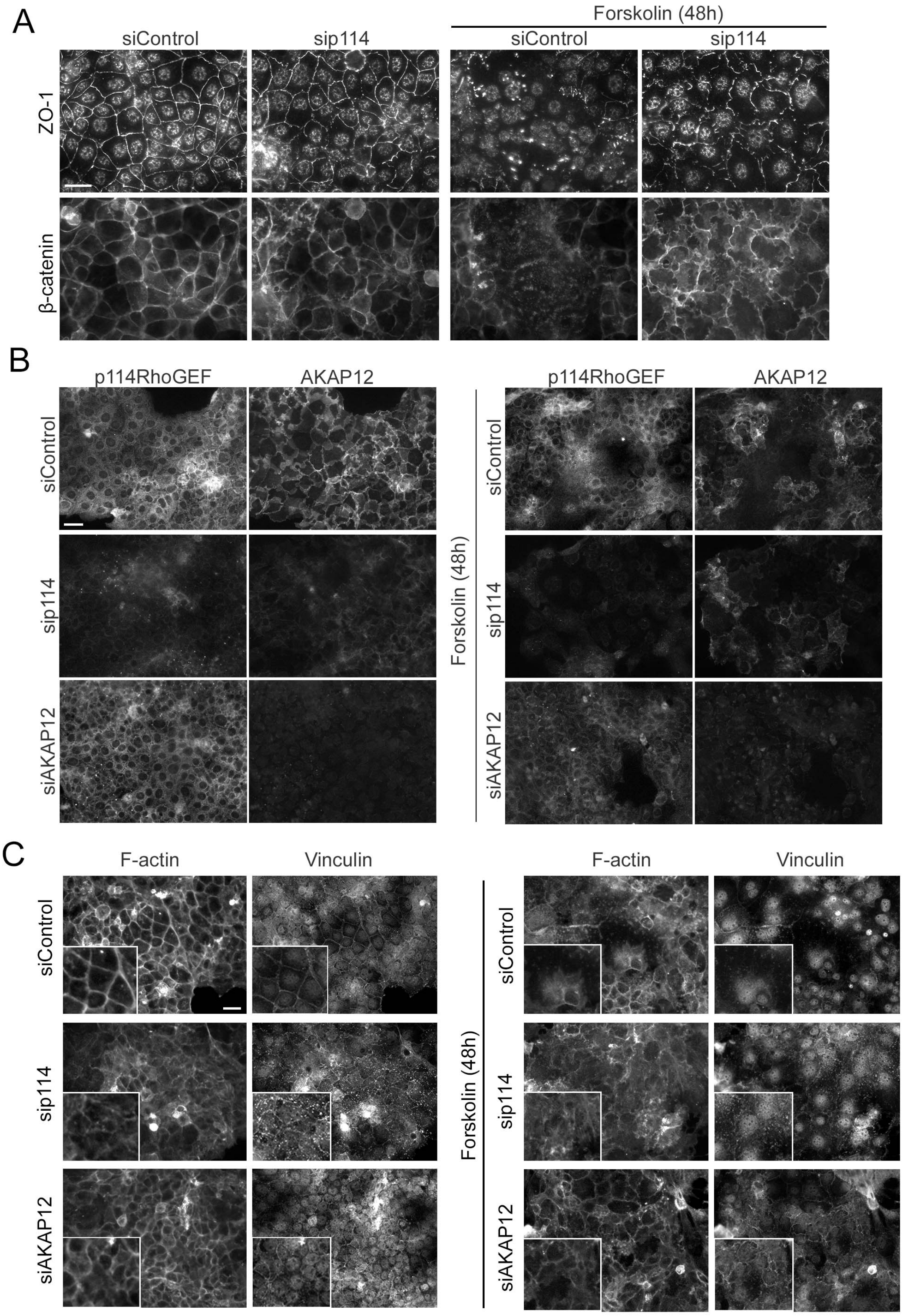


**Supplementary Figure 6. Inhibition of cell-cell fusion of BeWo cells.**
BeWo cells transfected with control, p114RhoGEF or AKAP12 siRNAs were treated with 100μM forskolin for 48h and then analysed by immunofluorescence for the junctional markers ZO-1 and β-catenin (**A**), p114RhoGEF and AKAP12 (**B**), or F-actin and Vinculin (**C**). Bars: 40μm.
